# Antisense inhibition of *accA* in *E. coli* suppressed *luxS* expression and increased antibiotic susceptibility

**DOI:** 10.1101/747980

**Authors:** Tatiana Hillman

## Abstract

Bacterial multiple drug resistance is a significant issue for the medical community. Gram-negative bacteria exhibit higher rates of multi-drug resistance, partly due to the impermeability of the Gram-negative bacterial cell wall and double-membrane cell envelope, which limits the internal accumulation of antibiotic agents. The outer lipopolysaccharide membrane regulates the transport of hydrophobic molecules, while the inner phospholipid membrane controls influx of hydrophilic particles. In *Escherichia coli*, the gene *accA* produces the acetyl-CoA carboxylase transferase enzyme required for catalyzing synthesis of fatty acids and phospholipids that compose the inner membrane. To increase antibiotic susceptibility and decrease growth, this study interrupted fatty acid synthesis and disrupted the composition of the inner membrane through inhibiting the gene *accA* with antisense RNA. This inhibition suppressed expression of *luxS*, a vital virulence factor that regulates cell growth, transfers intercellular quorum-sensing signals mediated by autoinducer-2, and is necessary for biofilm formation. Bacterial cells in which *accA* was inhibited also displayed a greater magnitude of antibiotic susceptibility. These findings confirm *accA* as a potent target for developing novel antibiotics such as antimicrobial gene therapies.

## INTRODUCTION

Multiple-drug resistant (MDR) bacteria, which are resistant to more than one antibiotic, present an enormous challenge for medical communities and organizations worldwide. For example, *Acinetobacter baumannii* is a highly contagious MDR gram-negative bacterium that inhabits hospitals and causes 64% of urinary tract infections associated with the use of catheters [1]. Many multiple drug resistance cases are caused by Gram-negative bacteria, which have a double-membrane cell envelope that provides partial protection from diffusible hydrophobic and hydrophilic compounds. This cell envelope consists of an inner membrane (IM) with a phospholipid bilayer, a periplasmic space, and an outer membrane with a lipopolysaccharide (LPS) region. LPS blocks the influx of hydrophobic molecules, while the IM constrains the internal transport of hydrophilic molecules, thereby limiting the accrual of antibiotics and enzyme inhibitors within the bacterial cell. Discovering antibiotic drugs that are effective despite this double-membrane structure is an arduous task [2].

The phospholipid (PL) bilayer of the Gram-negative cell envelope is produced through the fatty acid type II synthesis pathway (FASII) [2]. FASII proteins are expressed as cohesive and connected domains on a multi-polypeptide chain, and the acetyl-CoA carboxylase (ACCase) complex is an essential enzyme for the first-rate limiting step in fatty acid synthesis. ACCase consists of three subunits: a dimerized biotin-dependent carboxylase (BC), a biotin carboxyl carrier protein (BCCP), and an α2β2heterotetrameric carboxyltransferase (CT) [3, 43-46]. The genes encoding the ACCase complex are: *accA* and *accD*, which respectively encode the α and β subunits of CT; *accB* (BCCP); and *accC* (BC). Of those, *accB* and *accC* are transcribed simultaneously as the *accBC* mini operon [4].

In the first phase of FASII, carboxylation of acetyl-CoA by BCCP and CT produces malonyl-CoA. Specifically, the AccA subunit transfers a carboxyl delivered through BC-BCCP to CT, BC-BCCP positions a carboxyl taken from a glucose-derived pyruvate for CT, and CT orients the carboxyl to the acetyl-CoA, enabling the synthesis of malonyl-CoA (Fig 1). Subsequently, the malonyl-CoA is converted to malonyl-ACP and then combined with acetyl-CoA by the enzyme 3-oxoacyl-[acyl-carrier-protein] synthase 3 (FabH) in an ATP-dependent reaction to generate ketoacetyl-ACP [5]. After FASII is initiated, the CT enzyme catalyzes fatty acid elongation, adding two carbons per every cycle of the process (Fig 2).

**Figure 1.**
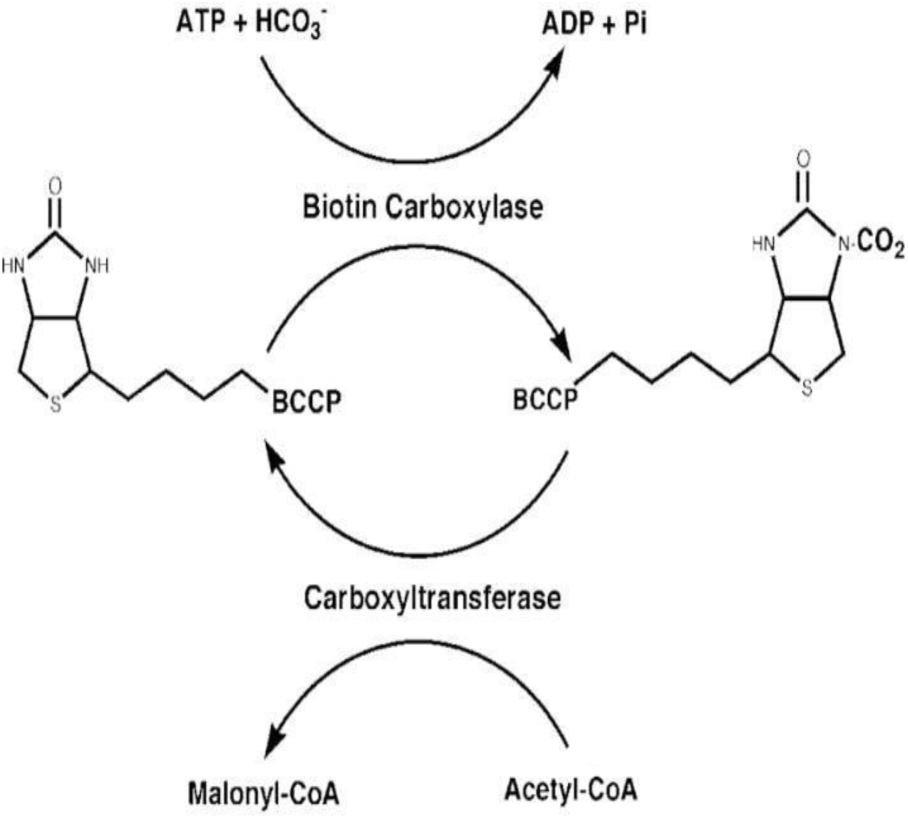
Formation of Malonyl-CoA. (*produced from reference 6 and originally published in the Journal of Biological Chemistry)*. The alpha and beta subunits of the AccABCD complex transfer a carboxyl group from pyruvate to acetyl-CoA, forming malonyl-CoA. The carboxylate moiety is carried by biotin covalently bound to BCCP through the action of a biotin carboxylase (BC)with ATP and carbon dioxide from bicarbonate as cofactors. In the second step of the reaction, a carboxyltransferase (CT) transfers carbon dioxide from carboxybiotin to a carboxyl group and then moves the carboxyl group to acetyl-CoA, producing malonyl-CoA.

**Figure 2.**
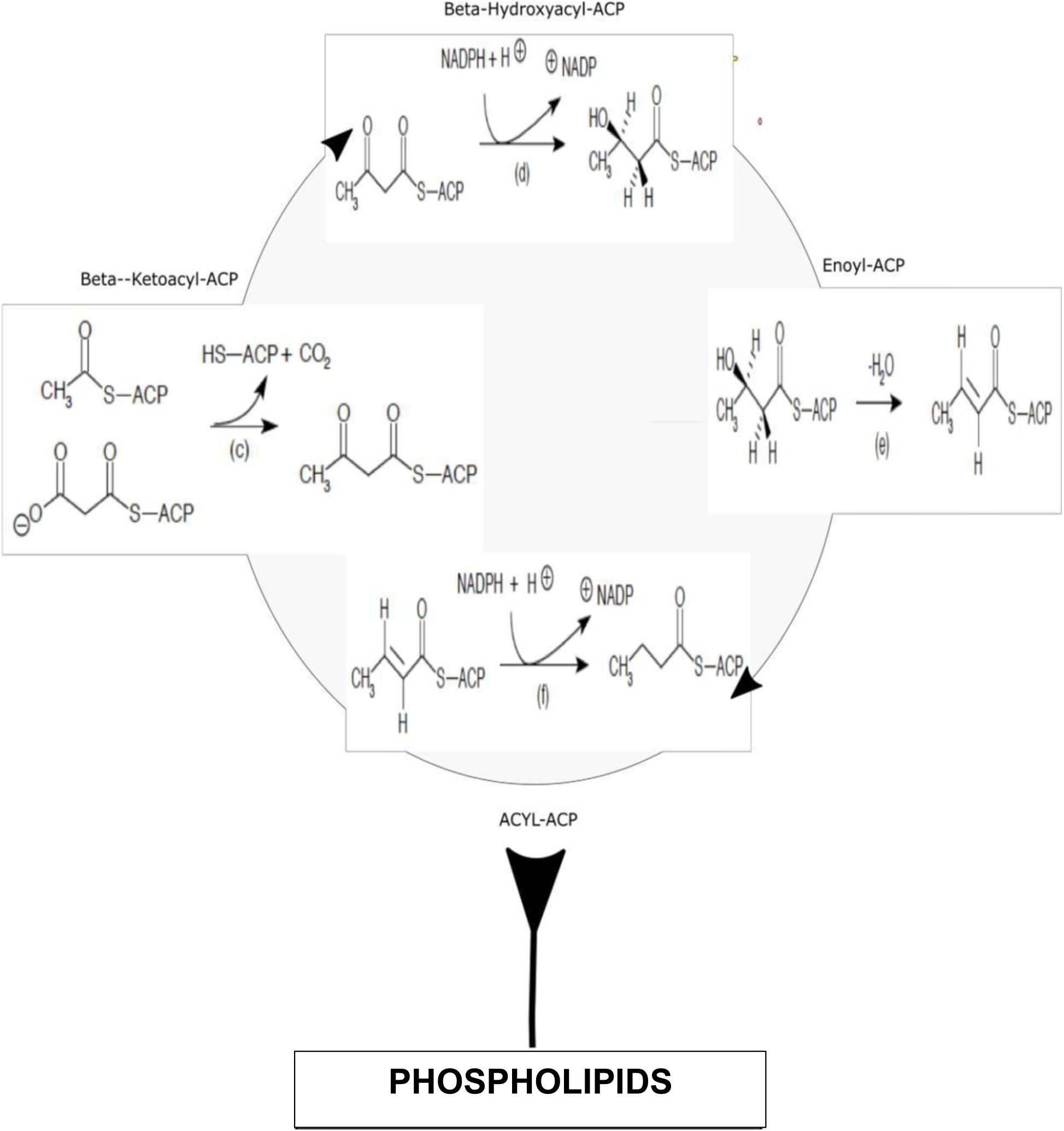
Long-Chain Fatty Acid Synthesis, An acyl-group, or ACP, is added to the malonyl-CoA and converted into ketoacyl-ACP by an added carbon dioxide. The Beta-ketoacyl-ACP is reduced and gains hydrogen into Beta-hydroxyacyl-ACP. The Beta-hydroxyacyl-ACP is dehydrated, losing a molecule of water, to alter into Enoyl-ACP, which forms Acyl-ACP by losing two hydrogen protons. Two carbons for FA elongation are added per cycle of FAS.

In addition to directly impacting membrane construction and being generally crucial for cell growth, bacterial fatty acid synthesis can significantly impact biofilm production, another element in antibiotic susceptibility and a major virulence factor. The extracellular biofilm matrix prevents the penetration of antibiotics and is mostly anaerobic, thus bacteria within a biofilm are more tolerant of antibiotics [7]. More research is needed to fully elucidate the link between the phospholipid membrane and biofilms [8]; however, Yao and Rock confirmed a significant connection between biofilm formation and enzymes for fatty acid synthesis in *Staphylococcus aureus* [9, 47]. A biofilm is characterized by tight junctional communities of bacterial cells that have an extracellular matrix covering allowing them to attach to a biotic or abiotic surface [10], and has been noted to feature a heightened presence of long-chain fatty acids that improves the orientation of fatty acids in the biofilm’s lipid bilayer, amplifies the links between acyl chains, and stabilizes the lipid bilayer [8]. This ultimately produces a more fluid membrane, and that fluidity causes the biofilm to expand, increasing the biofilm’s protective effect against external and environmental pressures.

However, the metabolic fluxes that affect bacterial biofilm formation are not yet fully elucidated in the literature, and the link between bacterial metabolism and virulence requires further explanation. Stokes et al. confirms that we are currently in the beginning stages of understanding the link between bacterial metabolism and antibiotic efficacy [27, 73-78]; a greater understanding of the interaction between bacterial metabolic potency and virulence is likewise needed [28]. Accordingly, the first study objective was to demonstrate the link between bacterial fatty acid metabolism and biofilm creation where the current study disrupted fatty acid synthesis in *Escherichia coli* through silencing the *accA* gene.

A second study objective was to ascertain possible interdependent activity between *accA* and *luxS*. LuxS is an autoinducer-2 (AI-2)-producing S-ribosyl homocysteine lyase protein also required for biofilm formation; a prior study in *Bacillus subtilis* demonstrated biofilm growth to be suppressed in the absence of vital nutrients (i.e. glucose) within cells [10]. Furthermore, Zuberi et al. applied CRISPRi technology to inhibit *luxS* in *E. coli* and found that biofilm formation as determined by crystal violet test depended upon its expression [12]. LuxS produces AI-2 from the precursor molecule 4,5-dihydroxy-2,3-pentadione (DPD). AI-2 is a released signaling molecule that induces quorum sense (QS) through bacterial cell-to-cell interactions; as such, it monitors metabolism, assists with biofilm formation, allows the transport of metabolite molecules, and transports regulatory signals [13, 48]. More specifically, as the biofilm broadens due to fatty acid incorporation, increased LuxS activity is needed to produce and release the AI-2 signals required for the continuum of quorum sense activities, thereby amplifying bacterial growth in proportional to the size of their extracellular matrix.

As noted, when Lux-S and AI-2 are less regulated and over-expressed, infectious bacteria can propagate in stressful environments with high acidity and salinity [39]. Kendall et al. demonstrated the ability of *S. pyogenes* to adapt to acidic conditions because of an upregulated Lux-S/AI-2 system [40]. The mutation of *luxS* in *Haemophilus influenzae* increases virulence and then causes middle ear infections [38]. The many facets of biofilm to tolerate antimicrobial therapy include many different types and subtypes of metabolic actions that regulate the bacterial response to different metabolic stimuli. The subtypes of metabolic changes include responses to pH, osmotic pressure, and the availability of nutrients [8]. Thus, LuxS and AccA can each affect bacterial biofilm formation. Finally, beyond investigating a connection between bacterial metabolism (glucose-*accA*-FASII) and virulence (i.e. AccA-LuxS activity), a third study objective also aimed to demonstrate the potential of *accA* as an advantageous target for novel antimicrobial gene therapies that ablate its expression.

### HYPOTHESES

To analyze the potential synergy between AccA and LuxS, *E. coli* (+)asRNA-*accA* transformants were exposed to glucose molecules, after which the expression of *luxS* was quantified by qPCR (*luxS*-qPCR). In addition, to test antibiotic susceptibility, tetracycline-, carbenicillin-, and chloramphenicol-containing disks were implanted in *E. coli* (+)asRNA*-accA* cell cultures and zones of inhibition measured. Two main hypotheses were tested in this study: 1) silencing *accA* and lessening glucose availability decreases *luxS* expression, which is essential to the LuxS/AI-2 QS system necessary for biofilm creation, and 2) inhibiting *accA* heightens the antibiotic susceptibility of Gram-negative bacteria because its suppression degrades the IM via FASII disruption.

## MATERIALS AND METHODS

### BACTERIAL CELL CULTURE AND RNA EXTRACTION

*Escherichia coli* (TOP10 strain) was cultured and plated on 25 mL agar plates. The bacterial cells were inoculated into Luria-Bertani broth enhanced with glucose at concentrations of 200 µM, 50 µM, and 0 µM (173 ± 72, N=21), replicated seven times (n=7) for quantification of bacterial dsDNA. Glucose was added to the *E. coli* LB liquid cultures with concentrations of 15 mM (High-glucose), 7.5 mM (Medium-glucose), 5 mM (Low-glucose), and 0 mM (control) (48±38, n=12). Antisense expression vectors were assembled using pHN1257-IPTG-PT plasmids. The antisense form of *accA* was inserted into pHN1257-IPTG-PT plasmids. The restriction enzymes of XhoI (upstream) and NcoI (downstream) were used to digest pHN1257 and the accA antisense gene insert for ligation (New England Biolabs, catalog no. R0146S and R0193S).

Cells transfected with recombinant pHN1257 plasmids for the asRNA of *accA* production were plated with the antibiotic kanamycin and incubated at 37 °C for 24 h. Cells grown on agar plates were inoculated into 4 mL of LB liquid media and evaluated for asRNA expression, classified as either (+) or (-)PTasRNA-accA. The total RNA was extracted for determining the RNA concentration of each bacterial sample. The total RNA was used to synthesize the cDNA for the qPCR analysis of *accA from E coli cells with* 15 mM, 7.5 mM, 5 mM, and 0 mM. The cDNA for *luxS* was synthesized for qPCR analysis, using primers specific for the *luxS* gene. The RNA was extracted, and the *luxS*-cDNA was synthesized from (+)-asRNA-*accA* expressing *E. coli* cells with 25 µM, 5 µM, and a control of glucose. About 25 total RNA extractions (335.4 ± 329.2, N=25) were completed with the Omega E.Z.N.A bacterial extraction kit.

### cDNA PREPARATION

The OneScript Reverse Transcriptase cDNA Synthesis kit by Applied Biological Materials Inc. was used according to the manufacturer’s instructions for cDNA preparation for both *accA* and *luxS* quantification. The total RNA was extracted from *E. coli* samples cultured with 15 mM, 7.5 mM, 5 mM, 25 µM, 5 µM, and 0 mM (control) of glucose. The primer/RNA mix consisted of 1 µg total RNA, 1 µL oligo(dt), and 1 µL of dNTP mix (10mM), 4 µL of 5x buffer, 1 µL of RTase enzyme, 0.5 µL of RNAse inhibitor, and 10.5 µL of water added up to 20 µL for the total reaction volume.

### PCR

PCR was performed using the Promega PCR MasterMix kit according to the manufacturer’s instructions with a reaction volume of 25 µL. Reaction-specific components consisted of upstream and downstream primers (0.5 µL) specific for the *accA* gene target from cDNA samples. The PCR reaction mixture consisted of 2X (5 µL) and 1X (2.5 µL) for upstream and downstream primers, less than 250 ng of DNA template, and nuclease-free water. The PCR amplification was completed in 40 cycles and PCR reactions were repeated in triplicate (n=3) (N=12, 219 ± 171). The thermocycler was set to a denaturation of a 2-minute initial denaturation step at 95°C. Other subsequent denaturation steps were between 30 seconds and 1 minute. The annealing step was optimized with annealing conditions by performing the reaction starting approximately 5°C below the calculated melting temperature of the primers and increasing the temperature in increments of 1°C to the annealing temperature. The annealing step was set at 30 seconds to 1 minute in 52°C. For the extension reaction with Taq polymerase, 1 minute was allowed for DNA to be amplified at 72°C with a final extension of 5 minutes at 72–74°C.

### AGAROSE GEL ELECTROPHORESIS AND PLASMID ASSEMBLY

PCR product quality in terms of primer dimers and contamination was determined through agarose gel electrophoresis. Subsequently, the PCR products and the PTasRNA expression vector (pHN1257) were digested with the restriction enzymes XhoI (upstream) and NcoI (downstream) (New England Biolabs, catalog no. R0146S and R0193S) (Fig. 1). The *accA* PCR products were ligated into the pHN1257 plasmid by mixing 1 µL of the DNA insert, 2 µL of plasmid, and 5 µL of the ligation mix (Takara Ligation Kit 6023), and then placing the tubes into a heat block at 90 °C for 15 min. Each microcentrifuge tube was placed in an incubator for 30 min at 37 °C.

### REAL-TIME PCR

Absolute quantification by real-time PCR was used to determine the expression levels of *accA* and *luxS. E. coli* pHN1257-(+)asRNA-*accA* transformants were cultured in media with glucose enhancement at concentrations of 15 mM, 7.5 mM, 5 mM, and 0 mM (N=14), respectively labeled as high-glucose (H-glucose), medium-glucose (M-glucose), low-glucose (L-glucose), and the control. The cDNA of *accA* synthesized from each treatment sample was diluted into 1:8 series dilutions to produce dilution curve tests for *accA*. Bacterial cells were transformed with pHN1257-(+)-asRNA-*accA* plasmids and *luxS* expression levels were analyzed by qPCR. Approximately, 25μM and 5μM of glucose was added to the (+)-asRNA-*accA* bacterial samples. The (+)asRNA-*accA* expressing cells were cultured without (-)glucose.

*E. coli* samples without glucose ((-)glucose) and without asRNA of *accA* ((-)asRNA-accA) were the control groups. RNA was extracted from each *E. coli* bacterial sample. About 1 µg of isolated RNA was reverse transcribed into cDNA for qPCR analysis. The cDNA was diluted into 1:10 dilution series to produce dilution curves for quantifying *luxS*. The value of gene copies for *accA* and *luxS* were measured using a qPCR amplification protocol (Roche LightCycler 480). The total reaction mixture volume included: 19.2µL of the primer pair mixture pipetted into a microcentrifuge tube with 240 µL of SensiMix SYBR green MasterMix, and with 28.8µL of nuclease-free water. About 12µL was pipetted from the SYBR green total reaction mixture into each well containing 8µL of the 1:8 dilution series for *accA*, the 1:10 dilution series for *luxS*, and the negative control.

### ANTIBIOTIC RESISTANCE TEST

Competent T7 *E. coli* cells were transformed with the pHN1257 inducible vector expressing antisense RNA to inhibit *accA* and plated on nutrient agar medium containing the antibiotic kanamycin. After transformant colony growth, 3-5 colonies were selected and cultured in LB broth for 18 to 24 h at 37 °C in a shaking incubator. A control group that did not express asRNA was likewise cultured in LB broth overnight at 37 °C. Subsequently, 100 µL of cultured broth was pipetted onto nutrient agar petri dishes (N=64) and spread with a swab to form a bacterial lawn; for pHN1257-asRNA-*accA* samples (n=8, 21±4, SEM=0.7, N=32). To each plate was added a disk containing 5 µL of the antibiotic for tetracycline (10 µg/mL), carbenicillin (100 µg/mL), and chloramphenicol (25 µg/mL). The bacterial cultures were then incubated for 18-24 h at 37 °C, after which the zone of inhibition for each disk (N=64) was measured in mm according to the Kirby-Bauer method and confirmed by the Clinical and Laboratory Standards Institute Disc diffusion supplemental tables for interpreting antibiotic resistance test results [84].

### STATISTICAL ANALYSIS

Descriptive analysis for the means and standard deviations were calculated using Microsoft Excel and GraphPad Prism to analyze nucleic acid production and the gene expression of *accA* after glucose addition to *E. coli* samples (N=41, M=365, SD=220). Standard curves for the absolute quantification of *accA* (n=4, N=16, M=48, SD=38) and *luxS* (N=12, n=4) were calculated using Microsoft Excel where linear regression coefficients were added to each dilution series curve. The descriptive statistics and an ANOVA analysis with Bonferroni and Holm results for the *p*-value of *luxS* gene copies (N=12, n=3, <u>M</u>=2.15×10^5^, <u>SD</u>=1.09×10^5^) were computed through GraphPad Prism. The antibiotic resistance test data (N=64, M=17.50, SD=2.00) were analyzed, and subjected to paired *t*-tests using OriginPro 2019B to calculate the p-value (*p*<0.05, paired *t*-test, t-score 3.6, df=31, 95% CI, [6, 20]). Figures illustrating the analyses were constructed with Sigma Plot (14th edition), Microsoft Excel, and GraphPad Prism (8th edition).

## RESULTS

### Glucose and *E*.*coli* Nucleic Acid Production

A paired t-test was used to analyze the level of bacterial yield of nucleic acid after treating *E. coli* samples with glucose. The DNA concentration was measured in ng/µL. In a paired t-test and when comparing 200 µM glucose bacterial samples to the control groups, the difference was statistically significant (*P*=0.038), with bacteria grown at 200 µM of glucose yielding substantially more dsDNA than the control sample (Figs 3-4). All conditions were significantly different from each other and for each bacterial sample. The 50μM of glucose in a bacterial sample highest yield was 566ng/μL, for 200μM, 117ng/μL of RNA was produced, and the control measured to 76 ng/μL of RNA. Bacterial samples with the highest amount of glucose produced higher levels of RNA transcription. The obtained total RNA concentrations were 1392, 797, 608, and 199 ng/µL for H-glucose (15 mM), M-glucose (7.5 mM), L-glucose (5 mM), and the control, respectively (N=25, M=335.4, SD=329.2)(Fig 3).

**Figure 3.**
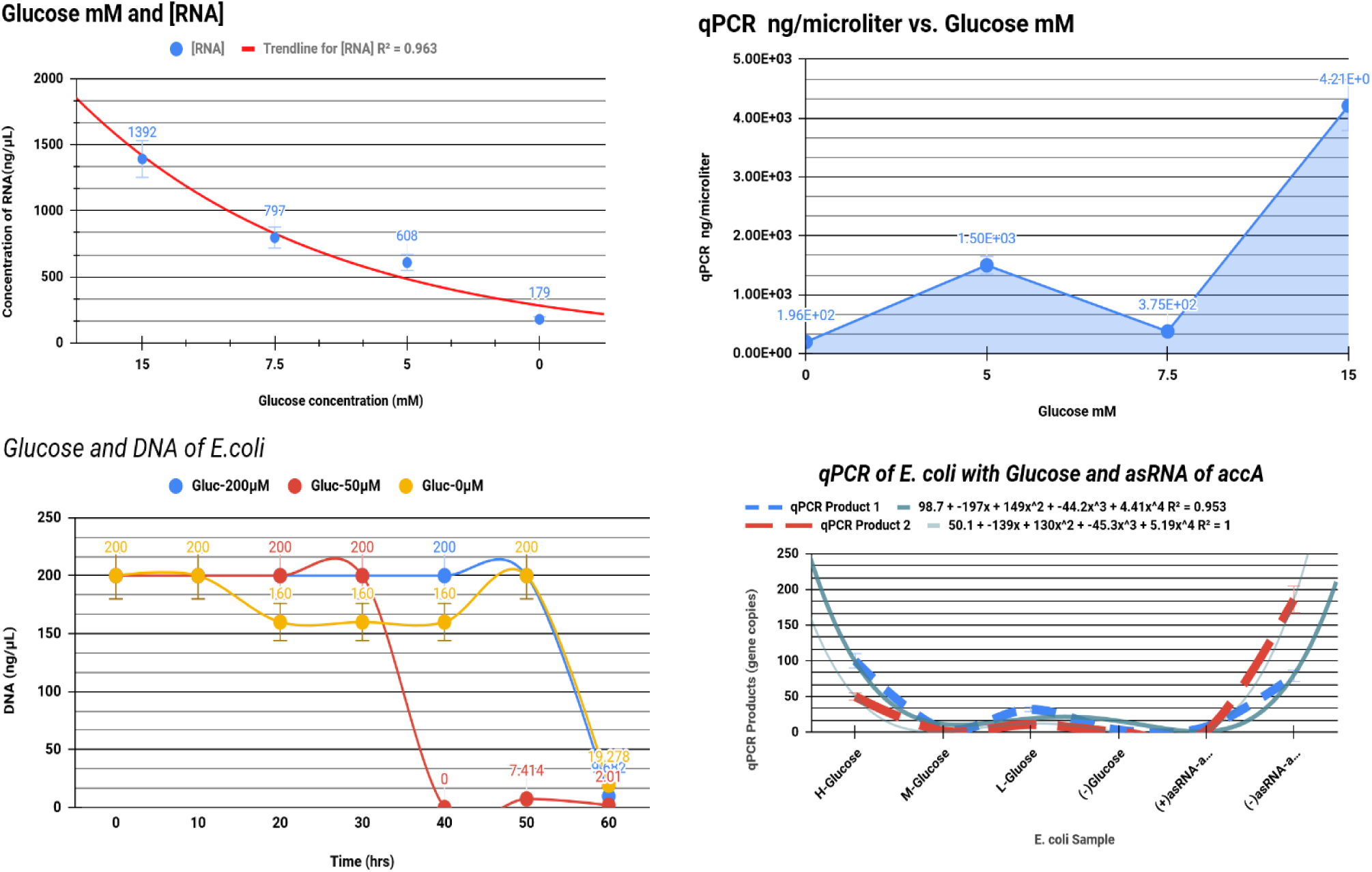
Influence of glucose concentration on RNA transcription and *accA*. Bacteria were cultured with added glucose at a range of concentrations (15 mM, 7.5 mM, 5 mM, or 0 mM). **A)** Direct proportional relationship of *E. coli* total RNA concentration and glucose. **B)** Transcription of *accA* was greatly increased at the highest glucose concentration tested (15 mM). **C)** DNA quantification based on OD260 for *E. coli* grown in glucose-enhanced media (200 µM, 50 µM, and 0 µM). **D)** qPCR quantification of accA gene copies, with the highest yield from H-glucose at 186 gene copies (n=4, 48±38).

**Figure 4.**
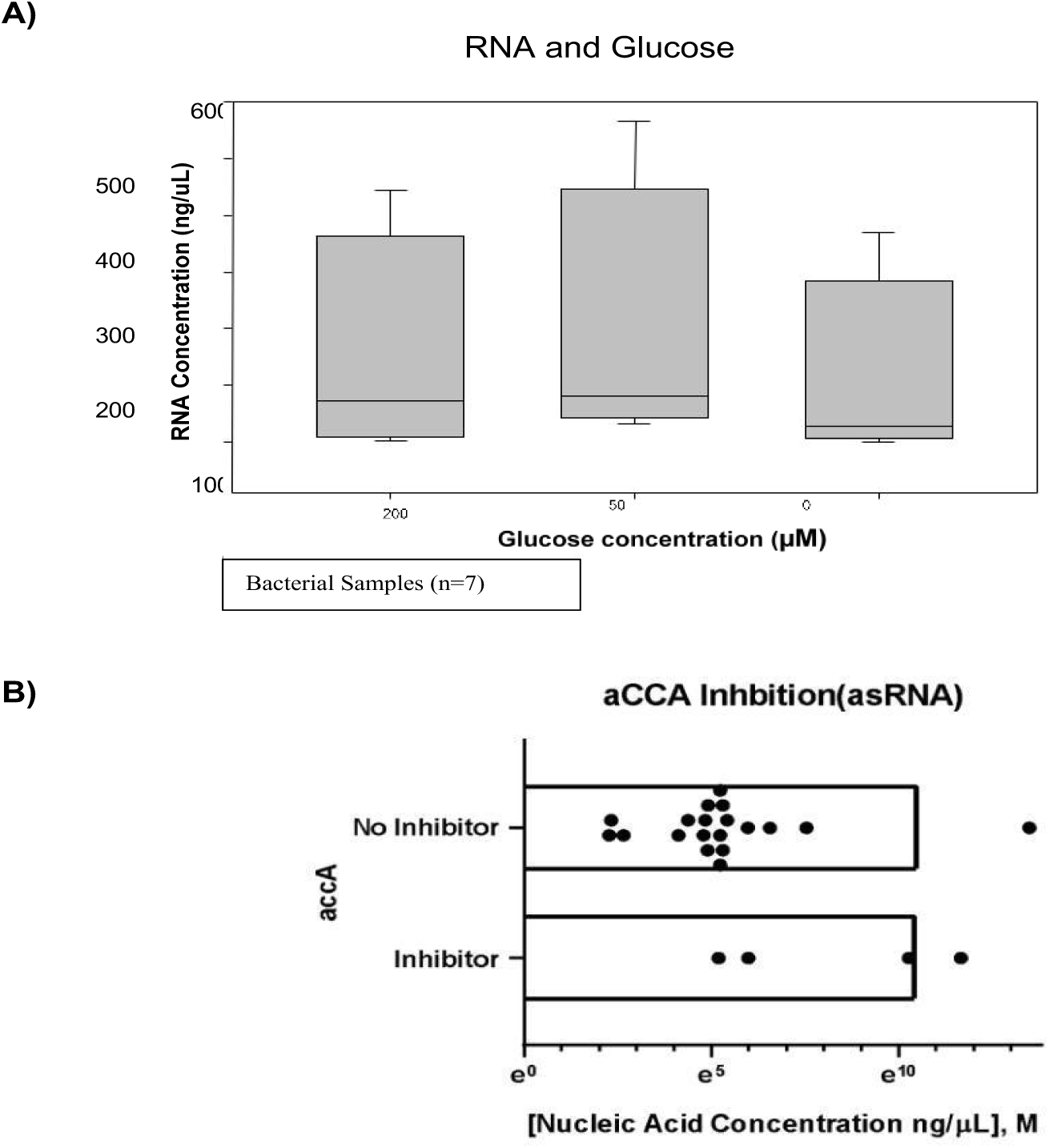
Relationship of RNA concentration to glucose concentration and effects of glucose and *accA* inhibition. **A)** *E. coli* (n=7) were cultured in media with 200 μM, 50 μM, or 0 μM of added glucose and total RNA quantified. **B)** Confirmation of *accA* transcription inhibition by asRNA. Samples expressing asRNA produced 16% (95% CI, [199, 1×10^6^]) of the total yield of nucleic acids, while those not inhibited yielded 83% (95% CI, [199, 1×10^6^]).

Similarly, the concentration of *accA* transcripts as determined by qPCR analysis were 4210, 375, 1500, and 196 ng/µL for H-glucose, M-glucose, L-glucose, and the control, respectively (Fig 3). Standard curves were used to perform absolute qPCR quantification of *accA* gene copies, with 11-186, 5-18, and 0-79 gene copies observed for H-glucose, M-glucose, and L-glucose/control samples, respectively (n=4, M=48, SD=38) (Fig 3). As glucose concentration increased, transcription in general and of *accA* was amplified. The residual plots of [RNA] versus [accA] were non-linear regressions with a non-constant variance that showed increased distance between the error and predicted values (Fig 5). There was a 147% difference between (+)asRNA-*accA* bacterial cells and the untransformed control groups, (95% CI [63, 421.69]). Suppression of *accA* was confirmed by qPCR, with a qPCR product of 63 ng/µL measured for (+)asRNA-accA *E. coli* cells (n=4) and 421.69 ng/µL for the (-)asRNA-accA cells. The obtained total RNA concentrations were 739.44 ng/µL for the control (-) asRNA and 279.28 ng/µL for bacteria transformed with pHN1257 plasmid expressing asRNA of *accA*, while the corresponding miRNA concentrations were 334.98 ng/µL and 240.90 ng/µL, respectively. Bacterial samples transformed with pHN1257 vectors for producing asRNA of *accA* showed a reduction in RNA transcription with less *accA* gene expression.

**Figure 5.**
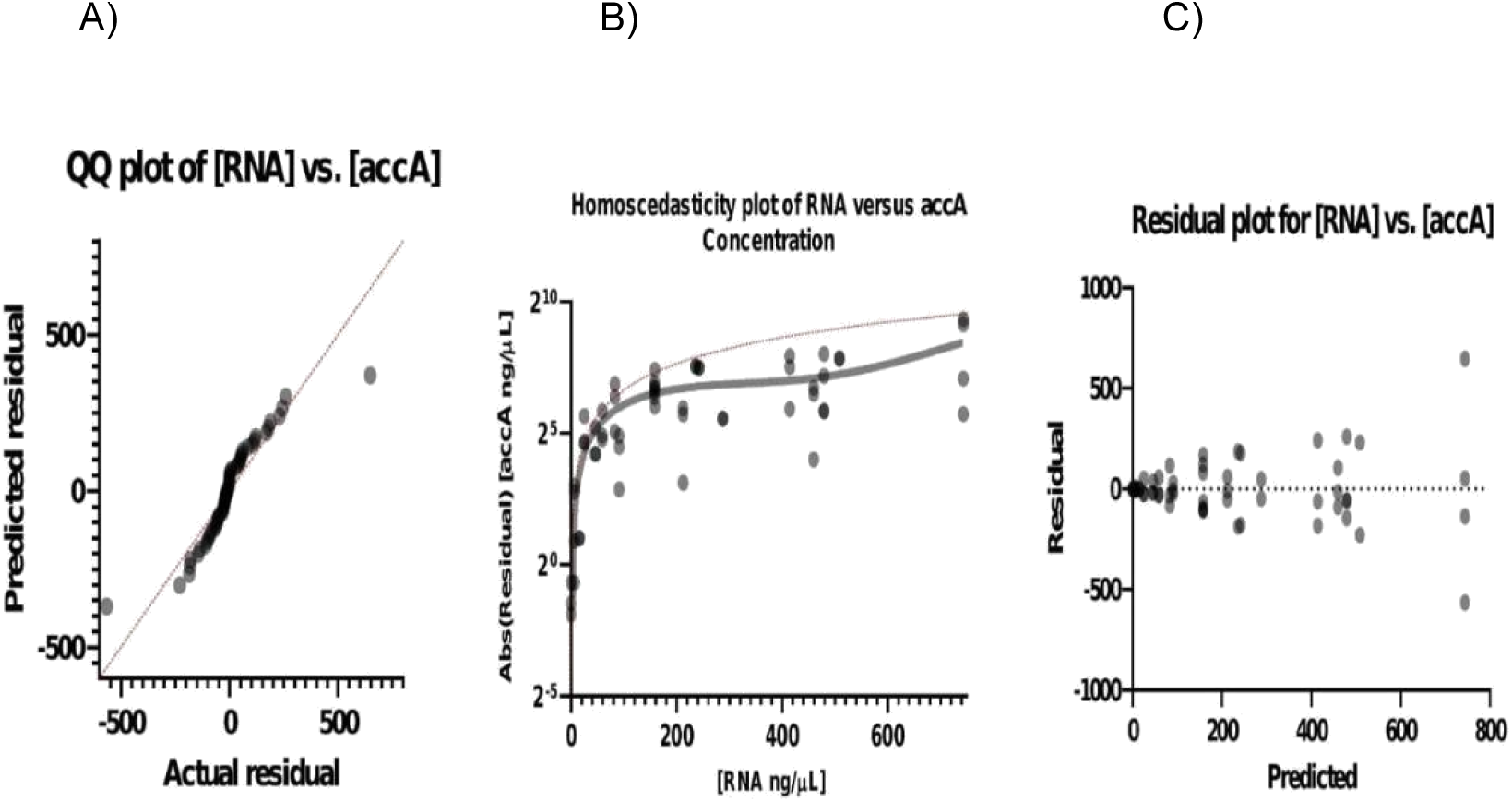
Predicted and actual residuals for RNA and *accA* concentrations. A) The QQ plot of RNA versus *accA* concentration exhibits means that are normally distributed. B) The homoscedasticity of [RNA] and [*accA*] is non-linear due to variance in error size. C) The residual plot resembles heteroscedasticity, with residuals increasing along with the predicted values.

### *luxS* and *accA*

A one-way ANOVA statistical test (Table 1) was performed to examine and to confirm this study’s hypothesis stating that less glucose and the inhibition of *accA* reduces *luxS* gene expression. To determine the effect of *accA* inhibition on *luxS* transcription, the number of l*uxS* gene copies were measured by qPCR after inhibiting *accA* and while altering glucose availability (25 µM, 5 µM, 0 µM). The concentration of *luxS* was measured in gene copy number values produced by the absolute quantification of the qPCR products. Because the independent variable of this method and approach used more than two dependent variables or conditions, a Bonferroni and Holm multiple comparison was implemented to determine which *E. coli* sample treatment group compared to the control groups were statistically significant at a *P*<.05 level (Table 2). All conditions were significantly different from each other except for the *E. coli* samples treated with 5 µM of glucose where the means were significantly lower than the standard deviation (Table 2). The control samples, the (-) glucose-(-) asRNA-*accA*, (n=3, 658,114 ± 483,499) and the 5μM glucose-(+) asRNA (n=3, 171571±204574) exhibited the highest amount of expression for *luxS*. The (-) glucose-(+) asRNA-*accA* sample produced a gene copy number of 199 (n=3, 277 ± 37) and 1×106 *luxS* copies for (-) glucose-(-) asRNA.

**Table 1.** Gene silencing *accA* lowered *luxS* levels in *E. coli* samples without glucose. The one-way ANOVA results showed the null hypothesis could be rejected, and the results supported the hypothesis for the silencing *accA* reducing *luxS* (F-value=5.0649, *p*=0.0296). The greatest difference of expression was observed between control (-) glucose-(-)asRNA and the (+)-asRNA-*accA* groups.

| source | sum of squares SS | degrees of freedom | mean square MS | F statistic | p-value |
| --- | --- | --- | --- | --- | --- |
| treatment | 1,028,447,998,008.9164 | 3 | 342,815,999,336.3055 | 5.0649 | 0.0296 |
| error | 541,473,744,144.0001 | 8 | 67,684,218,018.0000 |  |  |
| total | 1,569,921,742,152.9165 | 11 |  |  |  |

**Table 2.** Bonferroni and Holm result The *E. coli* samples treated with 5 µM of glucose were the only insignificant group of the treatment samples and conditions. The difference between the control (Treatment-A) versus the (+)asRNA-*accA* (Treatment-B) samples for the *luxS* qPCR products was significant (*P*=0.0291855). The control group against the 5μM-glucose-(+)asRNA-*accA* (Treatment-C) was insignificant (*P*=0.0660209). Control samples compared to the 25μM-glucose-(+)asRNA-*accA* (Treatment-D) *E. coli* samples were significantly different (*P*=0.0237462).

| treatment pair | Bonferro<br>ni<br>and Holm<br>T<br>T<br>-statistic | Bonferroni<br>p-value | Bonferroni<br>inference | Holm<br>p-value | Holm<br>inference |
| --- | --- | --- | --- | --- | --- |
| A vs B | 3.3739 | 0.0291855 | significant | 0.0291855 | * $p<0.05$ |
| A vs C | 2.8342 | 0.0660209 | insignificant | 0.0220070 | * $p<0.05$ |
| A vs D | 3.2403 | 0.0356193 | significant | 0.0237462 | * $p<0.05$ |

For the 25μM glucose-(+) asRNA and 5μM glucose (+), asRNA produced 39,810 and 316,228 gene copies of *luxS*, respectively. The difference between the control versus the (-)asRNA-*accA* samples for the *luxS* qPCR products was significant (*P*=0.0291855). The control group against the 5μM-glucose-(+)asRNA-*accA* was insignificant (*P*=0.0660209). Control samples compared to the 25μM-glucose-(+)asRNA-*accA E. coli* samples were significantly different (*P*=0.0237462). The asRNA inhibition of *accA* did lower *luxS* levels in the absence of supplemental glucose and each treatment was significantly different according to the one-way ANOVA test results (F-value=5.0649, *P*=0.0296) (Table 1), and the greatest difference of expression was observed between control (-) glucose-(-) asRNA and the (+)-asRNA-*accA* groups. The p-value corresponding to the *F*-statistic of one-way ANOVA is lower than 0.05, suggesting that one or more treatments were significantly different. Transcription of *luxS* also exhibited a proportional relation with *accA* levels (Fig 6).

**Figure 6.**
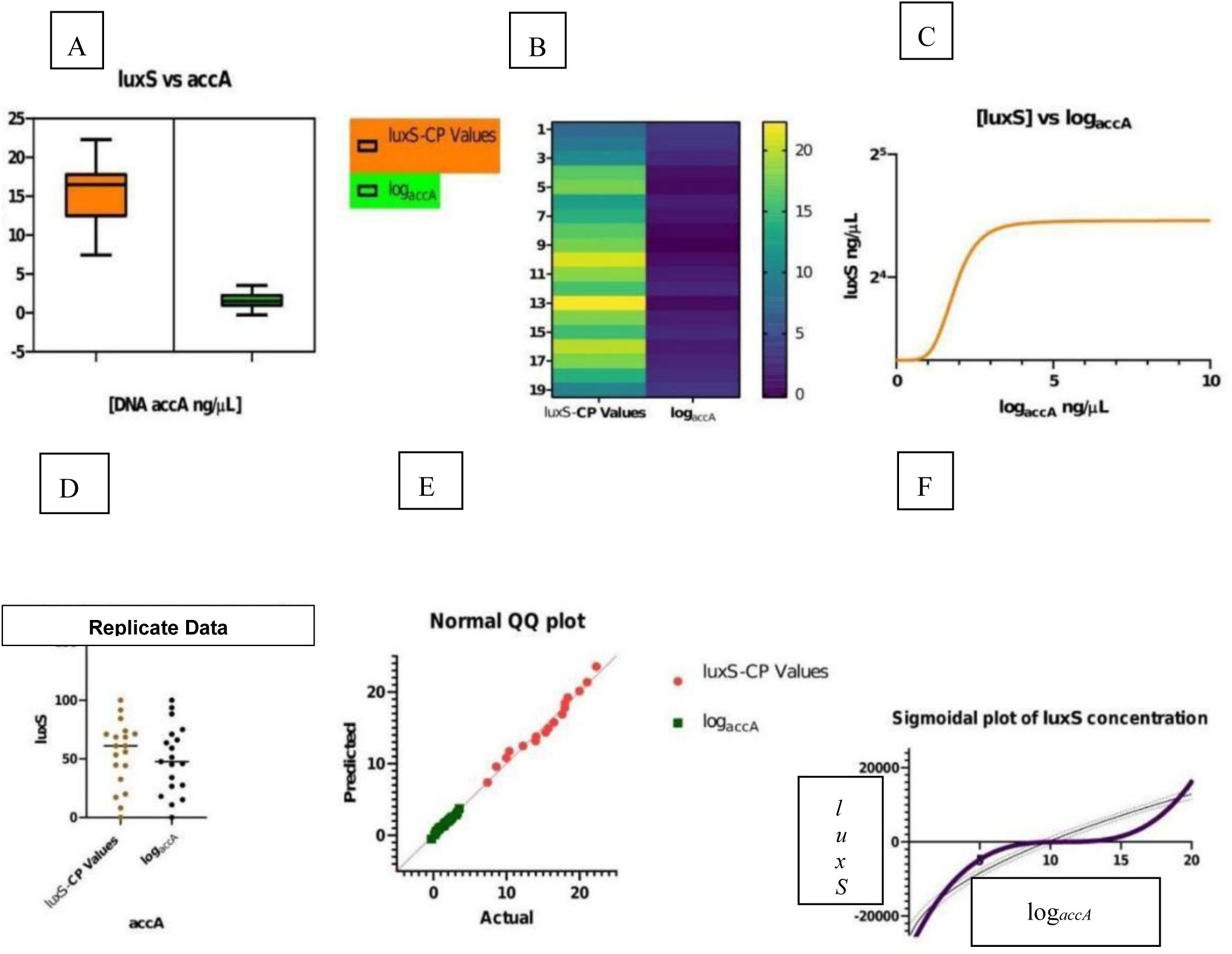
Effect of *accA* asRNA inhibition (log_*accA*_) on *luxS gene* expression (CP-Values). Transcription of *luxS* exhibits a proportional relation with *accA* levels. **A)** Inhibition of *accA* by asRNA resulted in tandem reduction of *luxS* expression. **B)** Heat map showing relationship of *accA* level and *luxS* Cp values. Higher CP values indicate less *luxS* expression; the highest Cp value of 20 at position 13 corresponds to a log_*accA*_ value near 0. **C)** Logarithmic relationship of *luxS* and log_*accA*._ As *accA* approaches 0, the expression of *luxS* decreases; as *accA* increases, *luxS* expression reaches equilibrium. **D)** Cp values for *luxS* parallel log*accA* data in most replicates. **E)** QQ plot showing normally distributed means for *luxS* Cp values and log_*accA*_. **F)** Sigmoidal plot showing a cooperative relation between log_*accA*_ and *luxS* CP.

### Antibiotic Resistance Tests

The level of antibiotic resistance was examined for *E. coli* cultures expressing asRNA for inhibiting *accA*. A paired t-test was used to support the second hypothesis for *E. coli* (+)-asRNA-*accA* samples exhibiting less resistance to antibiotics due to *accA* inhibition disrupting FAS, by damaging the IM, and then increasing antibiotic influx into mutant (+)-asRNA-*accA E. coli* cells. The antibiotic resistance was measured in mm after measuring the zone of inhibition for each antibiotic disk of tetracycline (10 µg/mL), carbenicillin (100 µg/mL), and chloramphenicol (25 µg/mL). All treatments were significantly different. *E. coli* samples exhibiting *accA* suppression (M=21, SD=4, SEM=0.7, N=32) were predominantly susceptible to all three antibiotics (Figs 7-8), with an average zone of inhibition of 20 mm. In contrast, the control samples (without (+)asRNA) were mainly resistant to the antibiotics, with an average ZOI of 6 mm. Ultimately, 59% of (+)asRNA-*accA* samples were sensitive to tetracycline, chloramphenicol, or carbenicillin showing larger zones of inhibition sizes. Another 28% displayed intermediate sensitivity, while 3% were resistant to antibiotics (13.75±10, SEM=1.8). Control samples exhibited higher prevalence of resistance, with 25%, 12.5%, and 53% of samples, respectively, demonstrating susceptibility, intermediate resistance, and resistance. Overall, the (+) asRNA-*accA* treatment group differed in sensitivity from the control group by 82% (95% CI, [6, 20]), and the control group differed in resistance from the treatment group by 77% (*P*=0.000554, paired *t*-test, t-score 3.6, df=31, 95% CI, [6, 20]).

**FIGURE 7.**
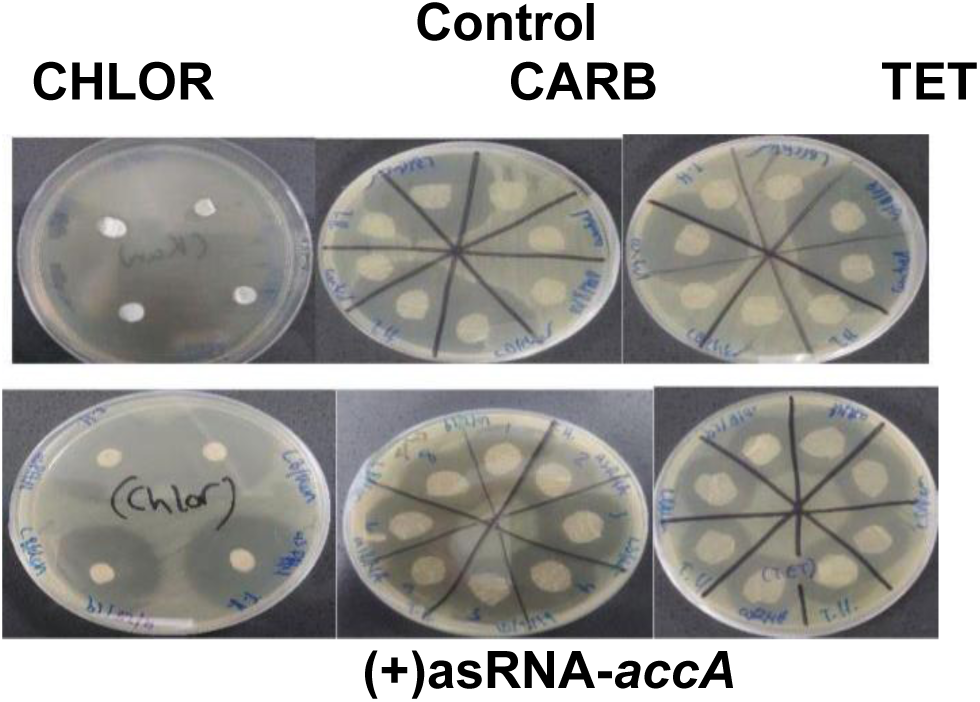
Antibiotic susceptibility testing. Bacteria transformed with pHN1257-(+)asRNA-*accA* showed greater susceptibility to antibiotic disks containing tetracycline (TET), chloramphenicol (CHLOR), and carbenicillin (CARB). Twenty-seven of the total 64 samples were susceptible; 13 demonstrated intermediate susceptibility; and 18 were resistant. The ratio of zone sizes for (+)asRNA-*accA* to (-)asRNA-*accA* was 3:2.

## DISCUSSION

### The Effect of Glucose on RNA Levels and *accA* Expression

To observe the effect of extracellular glucose on intracellular RNA transcription and *accA* expression, glucose was added to *Escherichia coli* liquid cultures, total RNA was extracted, and the RNA concentration determined. RNA was then isolated from the treated cultures to perform qPCR for quantification of *accA* expression (qPCR-*accA*). The *E*.*coli* samples with the highest concentration of glucose produced more RNA. Additionally, the increase in available carbon nutrients also increased transcription of *accA*, as reflected in the qPCR results. D-Glucose was added to *Escherichia coli* cultures positive for asRNA expression ((+)asRNA-accA) to affirm the interaction between glucose catabolism into carboxyls and *accA* gene expression. The augmentation of RNA transcription because of glucose supplementation occurred because increased carbon uptake induces intensified transcription of *accBC* and *accD* by controlling the flux of the tricarboxylic acid (TCA) cycle that augments the production of malonyl-CoA [15]. When *accA* expression was amplified, activity of the CT enzyme also surged, specifically transferring carboxyls to acetyl-CoA molecules for conversion into malonyl-CoA.

The bacterial sample treated with 15mM of glucose produced the highest concentration of RNA and *accA*. The biotin-dependent ACCase enzyme performs a crucial function for catalyzing fatty acid metabolic activity, completing two enzymatic reactions in two half-reaction steps. The first step, catalyzed by a biotin carboxylase (BC), is the carboxylation of the N1’ atom of a biotin vitamin cofactor; this depends on ATP and carbon dioxide derived from bicarbonate [5]. In the process, the biotin covalently bonds to a lysine side chain residue located in the biotin carboxyl carrier protein of the BCCP subunit. The biotin also interacts with the “thumb” of BCCP, a loop of eight amino acids that extends from the C-terminus domain [4] this fortifies the BCCP-biotin interaction. Then, in the second step, a carboxyl transferase (CT) transfers the carbon dioxide from the bonded carboxybiotin to a carboxyl group. Under increased glucose supplementation, excess carboxyl groups and ATP are available, promoting activity of the BCCP thumb and thus promoting the first carboxylation step in fatty acid synthesis. Glucose is also the source of the carboxyl groups transferred by the BCCP to CT. If a carboxyl is not present, biotin cannot attach to the CT subunit [16]. The control sample had a lower value of RNA and *accA* gene expression because without glucose providing carboxyl groups, then a biotin cannot bind to a CT for transferring carboxyls to acetyl-coA needed for catalyzing FAS.

In many bacterial species, carbohydrate metabolism also controls biofilm generation [19]. For example, increased glucose availability allows *Staphylococcus epidermidis* to upregulate transcription of genes specific for biofilm production during the stationary growth phase [20]. However, *Enterobacteriaceae* species suppress the creation of biofilm as glucose availability increases. Many other gram-positive bacteria also repress biofilm formation in the presence of glucose. *Enterococcus faecalis* can even have a strain-specific response, with some augmenting biofilm genesis when glucose is in excess, and others repressing biofilm construction while glucose is present [19]. Ultimately, the availability of nutrients determines the amount of biomass that accumulates within a biofilm, and glucose supplementation of the culture medium increases bacterial growth and total biomass.

### Confirmation of the Inhibition of *accA* by asRNA

Lipids are the central building blocks of the inner cytoplasmic membrane. Lipids maintain the structure of bacterial cells, increase survival, and increase growth, and allow the passage of hydrophobic units via increased membrane permeability. The lipid acyl chains that are produced during FASII construct more fluid and permeable bacteria cells. Bacterial cells located in the biofilm versus planktonic bacterial cells possess higher concentrations of fatty acids. The higher access of fatty acids increases tolerance to various temperatures, more stacking of bacterial cells, and more stable lipid bilayers. A heightened presence of long-chain fatty acids leads to the increased orientation of fatty acids in the lipid bilayer, amplifies the links between acyl chains, and stabilizes the lipid bilayer [8].

The phospholipid (PL) bilayer is produced by FASII [2] and the translation of *accA* into subunits AccA and AccD of the CT enzyme is needed for FASII catalysis in *E. coli*. Degrading the PL bilayer of the MDR-GNB inner membrane by inhibiting FASII is possible through silencing *accA* gene expression. To confirm the reduction of the gene *accA*, cells were transformed with pHN1257-IPTG-PT-asRNA vectors carrying the insert for asRNA inhibiting *accA*, after which total RNA was extracted and expression of *accA* quantified by qPCR. Specifically, *accA* was suppressed with interfering antisense RNAs (asRNAs), and its expression was quantified via real-time PCR (qPCR). Antisense RNAs were designed to target sequences flanking the ribosome-binding site (RBS) and the start codon of the 150-base pair (bp) *accA* mRNA, acting to block ribosomal detection and thereby inhibit translation [11].

The level of *accA* gene expression was greatly lessened to 63 ng/ µL, and the reduction of *accA* also lowered the RNA levels of *E. coli* samples. Lessening ACCase activity shortens the length of the acyl chain, which can increase or decrease in elongation. Bacterial cells, in response to low temperatures or toxicity, can change the ratios of their acyl chains through saturation to unsaturation, from cis to trans-double-bonded carbons, and from branched to unbranched stacks of acyl chains [8]. The fatty acids produced will also form a more fluid membrane, and this fluidity causes the biofilm to expand. As the biofilm broadens, increased LuxS activity is required to produce, and release AI-2 signals needed for regulating population growth via quorum sensing.

### qPCR Analysis of *luxS* expression Levels

In an intercellular process called quorum sense (QS) signaling, *E. coli* cells release AI-2 signaling molecules for detecting cell density and to regulate gene expression. In *E. coli*, the transporter and transmembrane complex called LsrABCD, an ATP-binding cassette importer, controls AI-2 uptake. LsrABCD contains transmembrane permease proteins, nucleotides for binding proteins, and a periplasmic solute to bind proteins. The repressor of LsrABCD and LsR is a kinase that is a part of the QS system of AI-2. Lsrk that is a kinase located in the cytoplasm, phosphorylates AI-2, and then the Lsrk attaches to the *lsr* repressor. Activated AI-2 suppresses the *lsr* repressor, termed LsrR, leading to increased release of AI-2 via autoinducer-activated expression of the LuxS protein. As the bacterial population increases, AI-2 fills the extracellular space [21, 49-58], for the detection of quorum sense signals where the bacterial cells can then alter gene expression and control the population density [22, 59-63]. Thus, quorum-sense activity monitors and regulates gene expression through a positive feedback loop triggered by the peripheral and internal accumulation of AI-2, which is ultimately dependent on the expression and translation of *luxS* (Fig 9).

The LuxS protein plays a significant role for generating infections [28]. LuxS/AI-2 controls expression of genes specific for affecting virulence and bacterial fitness as the biofilm of *S. pneumoniae* forms [36]. The accumulation of AI-2 is used by bacteria to infect more host cells, and AI-2 creates a more solid biofilm [37]. The activity of the flagella is amplified with higher levels of *luxS* mutations present in the lungs and bloodstream. The mutant *luxS* allows bacteria to infect the lungs and blood more than the wild-type strain [38]. LuxS produces the DPD precursor for AI-2 synthesis [28] where the purpose of LuxS and AI-2 includes regulating metabolism and monitoring quorum sense activity [28]. The *luxS* gene, when it is more expressed, causes the spread of virulent bacteria by increasing the motility of pathogenic bacteria.

LuxS is essential for cell growth, quorum sensing, and biofilm formation. To examine the link between FAS and virulent biofilm formation, *E. coli* cells were grown in glucose-supplemented or control media and transformed with pHN1257 plasmids expressing IPTG-PT-asRNA of *accA*. The hypothesis was supported by the results where the asRNA inhibition of *accA* did lower *luxS* levels in the absence of supplemental glucose (*F*=5.0649, *P*=0.0296). The greatest difference of expression was observed between the control (-) glucose-(-) asRNA sample and the (+) asRNA-*accA* samples. Glucose increased the rate of *luxS* transcription in cells versus cells without glucose. The control bacteria samples and the cells without asRNA of accA yielded 1×10^6^ and 199 copies of *luxS*, respectively.

Meanwhile, the heightened genetic activity at *luxS* observed upon glucose supplementation both with and without inhibition of *accA* supports the metabolic dependency of the LuxS/AI-2 system. For example, the results for the control group versus the 5μM-glucose-(+)asRNA-*accA* may have been insignificant (p=0.0660209) because according to Wang et al., adding 0.8% glucose to bacterial cultures and growth medium increases activity at the *luxS* promoter [17]. Wang et al. also found supplemental glucose increased the levels of AI-2 in culture [17]. However, Jesudhasan et al. (2010) demonstrated that when culturing *Salmonella typhimurium* in excess concentrations of glucose, higher levels of autoinducer-2 did not necessarily lead to increased *luxS* expression [18]. Hence, increasing glucose to 25 μM did not significantly amplify *luxS* gene expression. Overall, reducing the activity of the LuxS/AI-2 system, through suppressing *accA* and *luxS*, can reduce biofilm consistency and structure, which lessens bacterial virulence and reduces antibiotic resistance.

Inhibiting *accA* produced an incomplete fatty acid elongation cycle where the availability of FAs and acyl-groups decreased and lowered the fluidity of the cell membrane. Gram-negative bacterial transmembrane proteins are implanted in the hydrophobic interior of lipid bilayers, and FAS of lipids and phospholipids affect the composition of the cell membrane [23, 64-67]. The viscosity (thickness) or fluidity of the lipid bilayer is significant for the activity of membrane proteins. Membrane lipid bilayer thickness changes as the membrane fluidity and phases are altered for allocating proteins to different regions and phases of the bacterial cell membrane [24]. The varied configurations of lipids control the physicochemical properties of the cell membrane such as viscosity, rigidity, phase behavior, and membrane fluidity [25, 26, 68-72]. Unsaturated lipids have a double bond that creates a kink in lipids that prevent tight lipid-packing in the cell membrane and increases fluidity.

The integration of bacterial FAS catalysis by the ACCase-AccA-AccD of CT with *luxS* activity was evaluated where transcription of luxS exhibited a proportional relation with *accA* levels (Fig 6). Inhibiting *accA* reduced activity at the *luxS* promoter in the absence of supplemental glucose, due to reduced fatty acid synthesis. Jia et al. found the growth and metabolism of a *luxS* mutant strain of *Lactobacillus plantarum* decreased because there was a reduction of fatty acid metabolic synthesis, which degraded the cell membrane fluidity [29]. The quality of the biofilm, a major cause of virulence and a factor in antibiotic resistance, is largely determined by *luxS* expression. It may be inferred that disrupting FASII via *accA* inhibition can form an incomplete plasma membrane that has less membrane fluidity than needed for the LsrACDB transmembrane protein to operate, thereby impairing the influx of Al-2; this reduced internal AI-2 may have led to downregulation of *luxS* expression. Furthermore, an incomplete inner membrane can also permit an elevated influx of antibiotics into the cytoplasm of previously drug-resistant *E. coli*.

This study presented important results concerning the effect of *accA* inhibition along with glucose supplementation upon the expression of *luxS*, an essential component in bacterial biofilm formation, quorum sense signaling, and virulence. Inhibiting *accA* was observed to decrease *luxS* expression, with (+)asRNA-*accA* cell cultures producing 199 gene copies of *luxS* where the control yielded 1×10^6^ gene copies. This decrease of *luxS* is attributable to *accA* inhibition lessening FA production and decreasing cell membrane fluidity. Importantly, suppressing *luxS* can prevent biofilm formation and disrupt the LuxS/AI-2 QS system, thereby lessening the virulence of pathogenic bacteria. The hypothesis that silencing *accA* will lessen *luxS* gene expression was confirmed by the results (*F-value*=5.0649, *P*=0.0296). Antimicrobial gene therapies that target *accA* may provide a novel antimicrobial therapy with increased effectiveness against MDR-GNB because the *accA that* encodes AccA subunits of ACCase CT enzymes is conserved across many different bacterial species and can limit FASII catalysis.

### Antibiotic Resistance Tests

An antibiotic resistance test was performed to determine rather *E. coli* cultures expressing asRNA against *accA* were less resistant to antibiotics. The results supported the hypothesis that blocking *accA* lessens antibiotic resistance of *E. coli* bacterial cells (*P*=0.000554, paired *t*-test, t-score 3.6, df=31, 95% CI, [6, 20]). Bacteria exhibiting *accA* suppression (21±4, SEM=0.7, N=32) were predominantly susceptible to all three antibiotics such as to tetracycline (10 µg/mL), carbenicillin (100 µg/mL), and chloramphenicol (25 µg/mL) (Figs 8-9), with an average zone of inhibition of 20 mm. In contrast, the control samples (without (+)asRNA of *accA*) were mainly resistant to the antibiotics, with an average ZOI of 6 mm. Because bacterial metabolism and environmental stress play concrete roles in the outcomes of antimicrobial treatments [30],the inhibition of *accA* by asRNA disrupted fatty acid synthesis and elongation, which resulted in an unstable plasma membrane structure for the *E. coli* cells, which allowed a more facile transport of antibiotics across the inner membrane.

**FIGURE 8.**
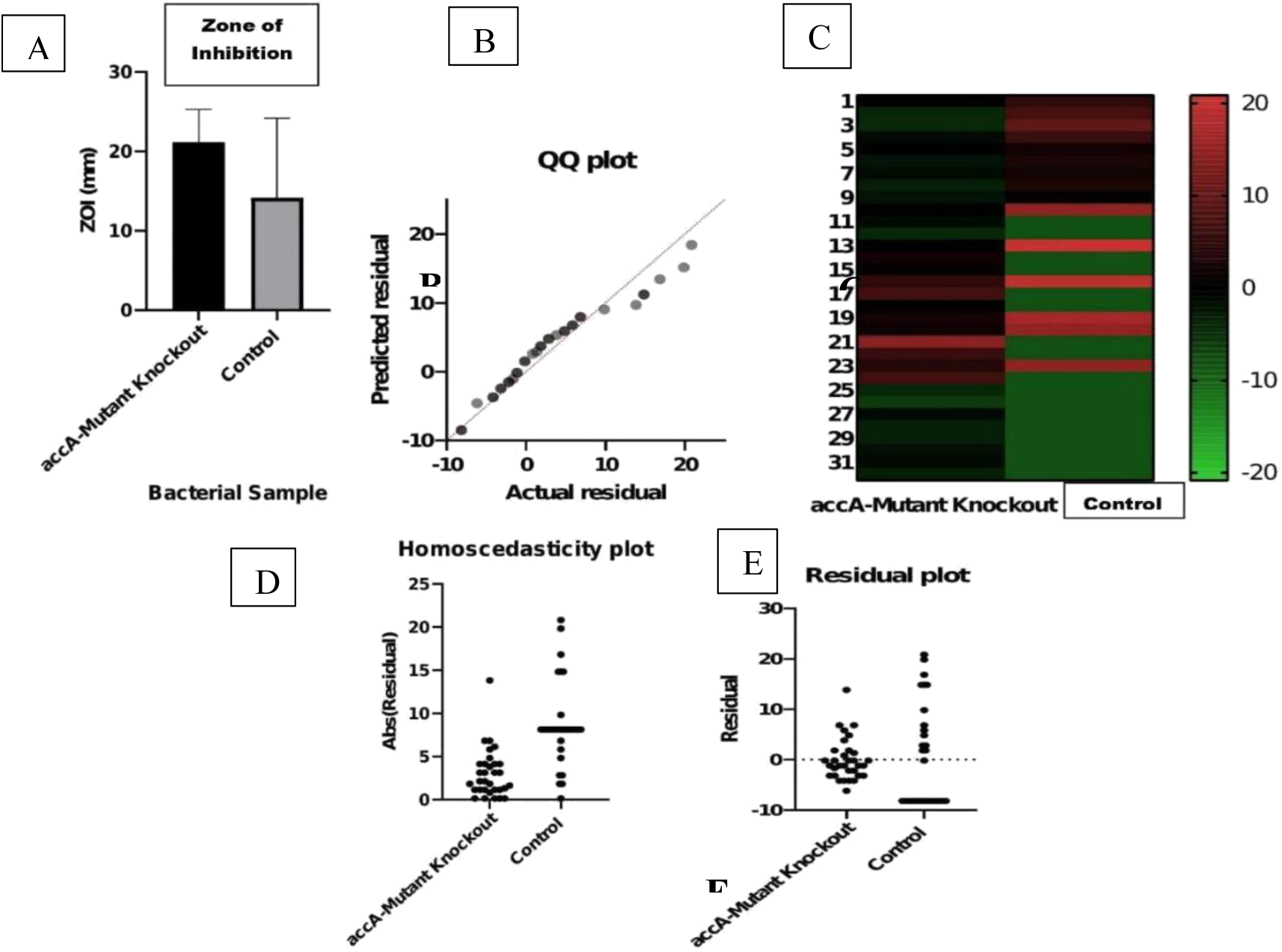
Comparison of the zones of inhibition. **A)** Zone of inhibition (ZOI) sizes differed by 60% (95% CI, [6, 20]) between asRNA-*accA* transformants and controls, with transformants having larger zones. **B)** QQ plot illustrating the normal distribution of data residuals for both transformants and controls. **C)** Heat map depicting relative zone of inhibition sizes (mm), with green indicating smaller ZOI and red for the larger zones. Controls tended to have smaller zones (greener). **D)** The homoscedasticity of residual data points is not maintained and is non-linear. **E)** The observed ZOI residual data points were different from predicted values for both transformants and controls, demonstrating a non-linear regression.

**Figure 9.**
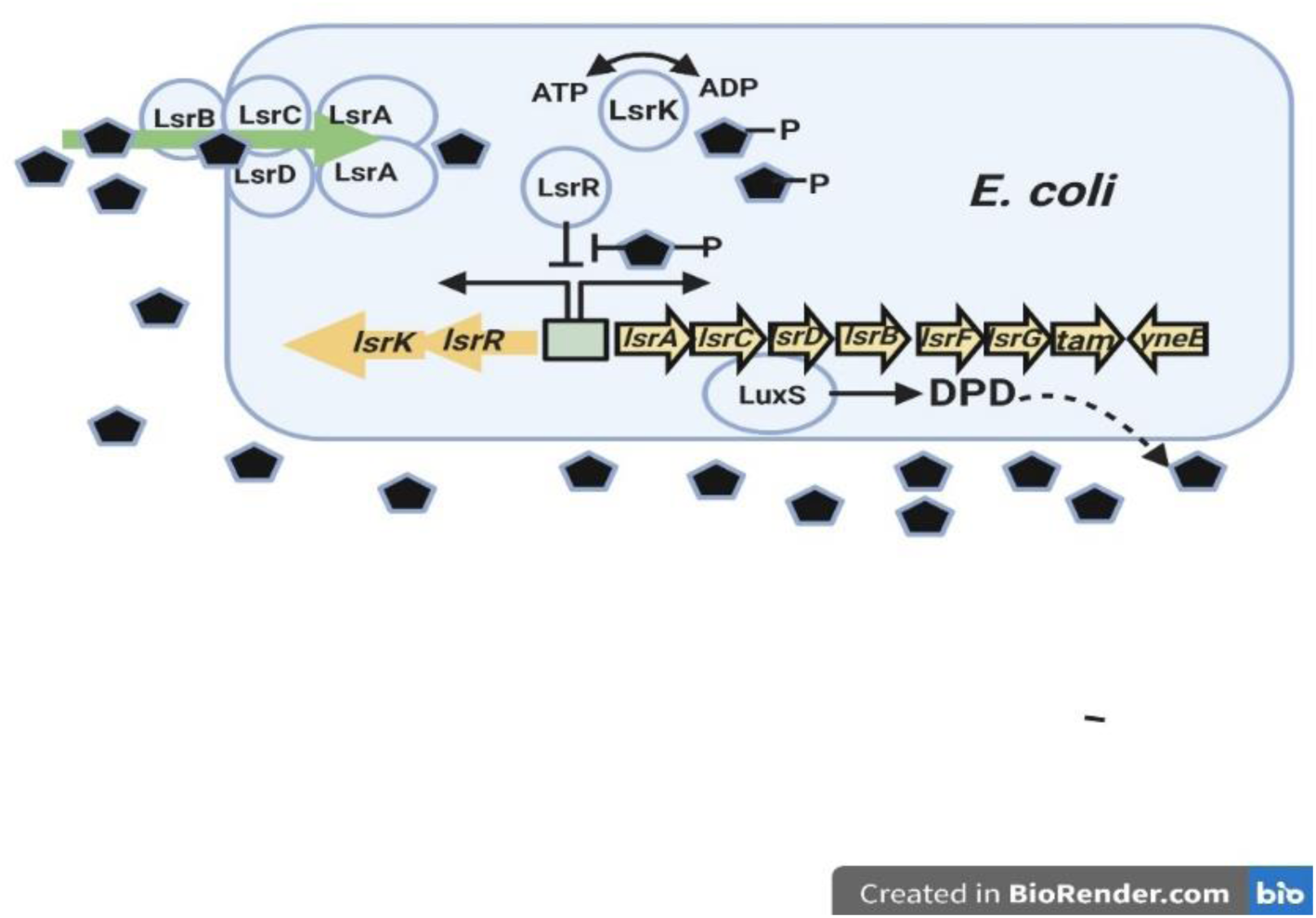
Regulatory mechanisms of the LsrR/phospho-AI-2 system in *E. coli*. (produced from reference *22*).The AI-2 (*black pentagons*) uptake repressor LsrR represses the *lsr* operon containing *lsrACDBFG* and the operon of *lsrRK*. AI-2 can re-enter the cell through the expression of LsrACDB. When the LsrK kinase phosphorylates an imported AI-2, the phosphorylated AI-2 binds to LsrR and terminates its repression of *lsr* transporter genes, inducing their expression. These regulatory steps trigger increased AI-2 uptake. (DPD=4,5-dihydroxy-2,3-pentanedione^) ***^ Created with BioRender.com

The potent metabolic state of a bacterial population can inhibit bacterial susceptibility to antibiotics [41, 79-83]; therefore, suppressing bacterial metabolism may decrease antibiotic resistance. Consistent with this expectation, the study findings showed *E. coli* cells expressing (+)asRNA-*accA*, in which the AccA subunit was eliminated from the ACCase-CT enzyme, were more susceptible to antibiotics. Furthermore, less FASII activity degrades the PL and IM of GNB, and this damaged GNB cell envelope allows a more facile influx of hydrophilic antibiotics. However, a few of the study findings were limited in that there was less access to hydrophilic antibiotics in the Kirby-Bauer antibiotic resistance tests. If more hydrophilic antibiotics were available, perhaps greater distinctions in ZOI and in the percent difference may have been demonstrated between the experimental and control groups. Nonetheless, the antibiotic resistance tests maintained and exhibited higher rates of susceptibility for (+)-asRNA-*accA* cell cultures where the inhibition of *accA* disrupted fatty acid production that damaged the IM structure allowing a greater influx of antibiotics.

In addition to their functional importance, the subunits of the ACCase enzyme complex are favorable targets for antibiotic design because bacterial ACCs are immensely different from mammalian and more specifically human ACC isoforms. Mammals depend on the isoform termed ACC2 or ACCβ; the two human isoforms ACC1 and ACC2 have 73% amino acid identity with ACCβ from mammals [5]. Additionally, eukaryotic cells have a BC domain with many partitioned and segmented residues; in particular, eukaryotic isoforms have 550 residues between the BC and AB domains, as compared to the 450 residues of bacterial BC units. An antimicrobial gene therapeutic designed to inhibit the expression of a bacterial ACCase enzyme thus cannot interfere or interact with a mammalian ACCase enzyme.

Notably, FASII-associated antibiotic adaptation involves robust bacterial proliferation and incorporation of host fatty acids salvaged from their environment. This may limit the potential of ACCs as metabolic targets for antimicrobial drug development. However, only *Lactobacillales* species are known to use exogenous fatty acids to completely bypass FASII inhibition [9]; in those bacteria, FASII shuts down entirely in the presence of exogenous fatty acids, allowing phospholipids to be solely synthesized from exogenous fatty acids. Inhibition of FASII cannot be bypassed in other bacteria because FASII is only partially downregulated in the presence of exogenous fatty acids or is required for synthesis of essential metabolites such as β-hydroxyacyl-ACP [9].

## CONCLUSION

Multidrug-resistant Gram-negative bacteria have created an intense challenge for clinicians treating infections of ill patients in intensive care units [26]; however, only in recent years has GNB become resistant to all common categories of antimicrobials [26]. During the past 30 years, the formulation and discovery of potent antibiotics have dwindled [31]. The higher rate of multiple resistance among GNB is partially attributable to their double-membrane cell envelope, which is comprised of an outer membrane consisting of LPS with endotoxins, a peptidoglycan cell wall, and an inner phospholipid membrane. The outer membrane can also restrict the penetration of antibiotics into GNB [32]. Many new and novel antibiotics have been produced in recent years to combat Gram-positive bacteria [33]; however, administration of these novel antibiotics has been limited to decrease further occurrences of MDR. Combined with the rise of MDR-GNB, this limitation has forced the administration of older antibiotics such as polymyxins B and E [34]. Antimicrobial peptides (AMPs) have been considered as an alternative novel approach due to their degrading bacterial structure through direct electrostatic interactions with microbial membranes [35]; however, many AMPs have failed to achieve success in clinical studies because they are also extremely toxic to mammalian cells.

Hall et al. concluded that the cause of antibiotic resistance or antibiotic sensitivity of biofilm originates from a genetic source [42]; therefore, perhaps combating MDR-GNB requires a gene-based antimicrobial approach as shown in the current study. This study’s findings of the interaction between fatty acid metabolism and virulence may serve as a first step to inform understanding of metabolic-dependent pathogenesis in bacteria and the findings confirm *accA* as a favorable and potential target for antimicrobial gene therapy development. The hypotheses were supported by the results where inhibiting *accA* reduced the *luxS* responsible for virulent biofilm growth and decreased antibiotic resistance. Building on this work could ultimately lead to future research that silences *accA* with small interfering RNAs or clustered regularly interspaced short palindromic repeats such as CRISPR-Cas9 nucleases. Future studies could focus on the application and delivery of siRNAs or CRISPR-Cas9 for silencing *accA* as an antibiotic gene therapy for MDR-GNB infections.

## ACKNOWLEDGMENTS

Special thanks are given to the staff from TheLAB in Los Angeles, CA. Thank you for your team. They were always available to answer questions and provide support.

